# Neuronal, Affective, and Sensory Correlates of Targeted Helping Behavior in Male and Female Sprague Dawley Rats

**DOI:** 10.1101/2022.08.17.503412

**Authors:** Stewart S. Cox, Brogan J. Brown, Samuel K Woods, Samantha J. Brown, Angela M. Kearns, Carmela M. Reichel

## Abstract

Empathy is an innate ability to understand the emotional states of others along with the motivation to improve it. It has evolved over time into highly complex behaviors, the basis of which can be described using the Perception Action Model (PAM), where shared affect promotes an action that eliminates the distress of both the passive “Target” and, by extension, the active “Observer.” There are myriad biological variables that may modulate empathic behavior, including sex, sensory modalities, and neural activity. In the following studies, using our lab’s model of social contact-independent targeted helping, we first tested whether sex differences exist in helping behavior. Next, we explored sex differences in sensory and affective signaling, including the impact of direct visualization of a distressed conspecific and the type of ultrasonic vocalizations (USV) made between animal pairs during the task. Finally, we examined the neural activity of multiple cortical and subcortical regions of interest across time during targeted helping between males and females. We show both sexes exhibit similar helping behavior, but sensory and affective signaling differs between sexes. Further, changes in neural activity exhibited distinct sex-specific patterns across time. Our results indicate sex differences are not a ubiquitous presence in targeted helping. Instead, it is likely sex differences may be a convergent phenomenon in which the behavior is similar, but the underlying biological mechanisms are distinct. These results lay the groundwork for future studies to explore the similarities and differences that drive empathic behavior in both males and females.

## INTRODUCTION

Empathy is a complex suite of behaviors that works to convey an understanding of the affective states of others. The ability to generate shared affect can have myriad benefits, including group cooperation, reproduction, and survival^1-2^. Empathic processes, therefore, help to inform interpersonal relationships, as well as guide complex social norms^3-4^. A prominent theoretical framework for understanding empathy is the Perception Action Model (PAM)^1-2, 5-6^. Complex empathic behaviors are built from and reliant on more simplistic ones, similar to a Russian nesting doll. At the core of these complex behaviors is the PAM^3,6^. According to the PAM, affective transfer occurs between a distressed animal (or “Target”) and a conspecific viewing the Target’s distress (known as the “Observer”), creating a shared affective state^7-8^. The Observer must regulate their new state to perform an action (e.g., consolation or aid) to reduce the distress of the Target, and, through a second emotional transfer, themselves^3,9^.

Numerous biological determinants are believed to contribute to the complexity of empathic behavior, including sex. While clinical research has historically shown women have consistently higher levels of empathic concern compared to men^10-11^, these conclusions have recently been called into question^12^. The few rodent studies using females also have conflicting conclusions on the importance of sex in empathic behavior^7, 13-14^. An exploration of the impact sex plays in empathic behavior is therefore necessary to advance the understanding of the field.

According to the PAM, empathic behaviors like targeted helping stem from an affective transfer between conspecifics. Sensory cues are critical for understanding the state of another and can modulate prosocial behaviors, but the mechanism behind this affective transfer is poorly understood and likely multimodal. Research suggests that rodent prosocial behaviors, such as social learning, may require the availability of visual cues for the best facilitation of learning^15^. As it more precisely relates to empathic behaviors, rodents have the ability to recognize the distress of another, at least in part, through visual cues^16^. Vicarious freezing in an emotional contagion paradigm was attenuated if an Observer’s view of the Target was obstructed by an opaque partition^17^. More recently, it was demonstrated that observational pain contagion in mice requires the image-forming visual system^18^. We therefore examined the role direct visualization of the distressed conspecific had on Observers during targeted helping in both males and females.

However, it is unlikely that the visual system is the only sensory modality critical in the transfer of emotional information between Observer and Target^19-20^. Increasing attention has been given to ultrasonic vocalizations (USV) as a proxy for understanding a rodent’s affective state. Although there is still a dearth of evidence regarding the behavioral consequence or specificity of USV in adult rats, they can broadly be categorized into two groups by frequency. Low frequency USV, often called 22 kilohertz (kHz) USV, with a range of 18-35 kHz, are emitted in the presence of aversive stimuli, such as a predator or predator odor^21^ and other stressors, like inescapable foot shock^22^. It is hypothesized that low frequency USV serve as alarm calls or indicate an aversive affective state. High frequency USV, also known as 50-kHz USV, fall within the range of >35 kHz and are emitted in prosocial situations like social exploratory activity, mating behavior, and other positive affective states^23-25^. The affective valence of rats during empathic behavior is not currently known^26-27^, but some evidence points to USV as critical for evoking a prosocial state and promoting helping behavior^7^. In the following experiments, USV of both the Target and Observer were recorded during multiple timepoints throughout targeted helping in males and females to understand their respective affective states.

A large library of clinical fMRI studies exists evaluating numerous substrates and their role in empathy. Brain regions that have been correlated with aspects of empathy include those involved in emotional salience and interoceptive valence, specifically the amygdala and insula^28-32^,as well as substrates necessary for perspective-taking, motivation, and cognition, like the prefrontal (PFC), anterior cingulate (ACC), and orbitofrontal (OFC) cortices^3, 33-37^. More causational and region-specific research using rats and mice are beginning to corroborate some of these imaging studies^8, 38-40^. There has been recent research exploring brain-wide neural activity during helping behavior in male rats^41^, but no work to date has examined sex differences in those regions. In order to examine the neural sex differences of prosocial behavior, activity of cortical and subcortical regions of interest was evaluated through immunohistochemical analysis of the immediate early gene *c-fos* across time and between sex during targeted helping.

In the following experiments, we used a model of social contact-independent targeted helping, in which Targets were placed in a compartment of the apparatus filled with 100 mm of water. Observers were placed on a dry platform with access to a chain that opened an automated door that, when opened, released the Target into a dry compartment separate from the Observer^42^. We used this model to investigate the behavioral, affective, sensory, and neural differences in targeted helping between males and females across three timepoints; early acquisition (EA), as an indication of an initial helping response; late acquisition (LA) to examine if additional experience with releasing a familiar conspecific modulates helping behavior as it does social interaction^43^; and reversal (Rev), to see if previous experience plays an equal role in helping behavior in males and females^42,44^ (see **Figure 1** for schematic representation of experiments and the experimental timeline). We conclude that sex differences are minimal in targeted helping behavior and can be attributed to a difference in task acquisition in a subset of animals. However, while the behavioral outcome is similar, it is likely that targeted helping may have sex-specific neural and affective mechanisms. A more complete evaluation of the sensory, affective, and neural components of targeted helping allows a better understanding of the distinct processes of complex prosocial behaviors in both males and females.

**Figure 1.**
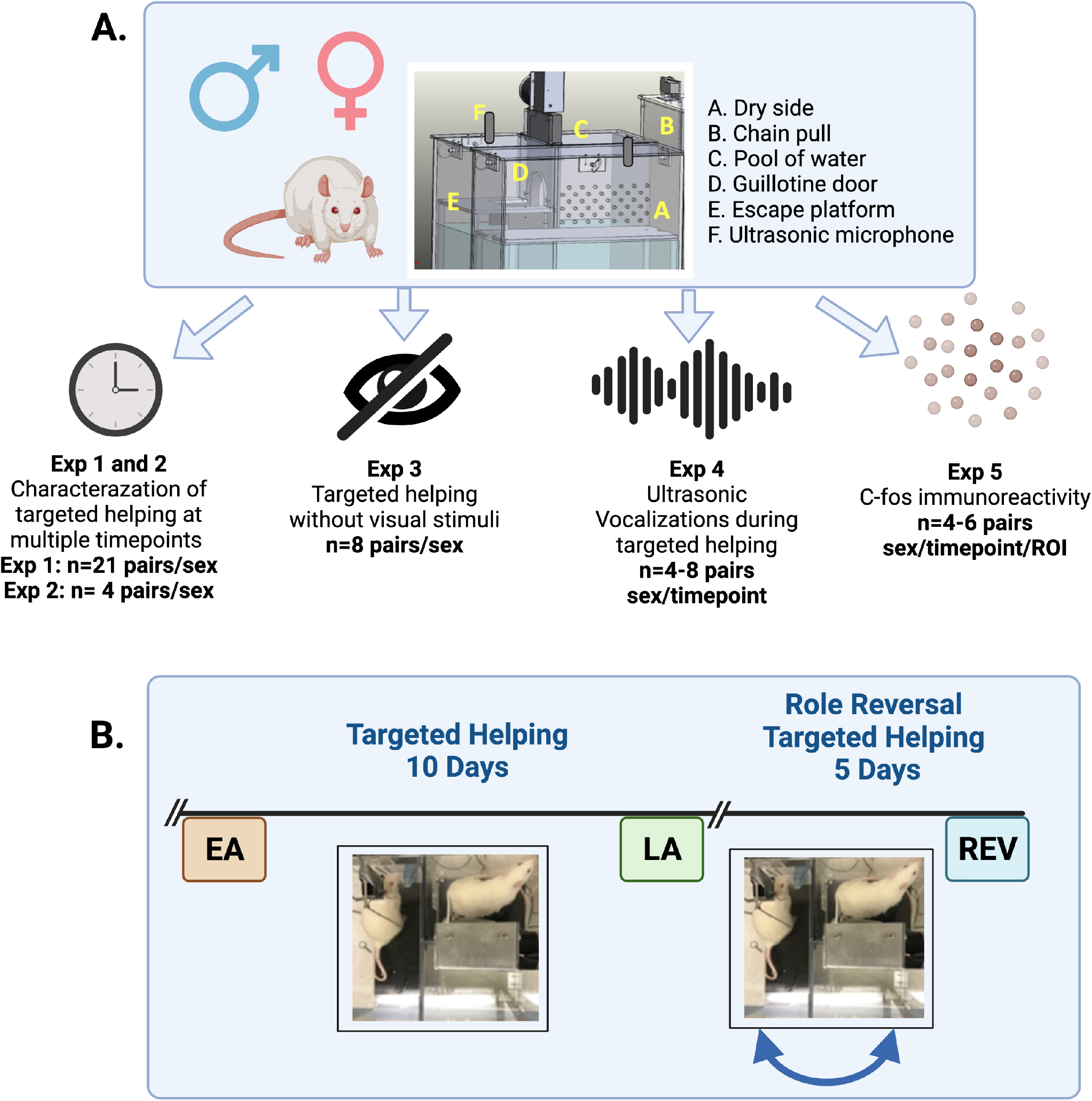
Schematic of Variables Tested and Experimental Timeline. A total of 33 pairs of rats/sex were used in Experiments 1-5. **A)** Size-matched, same-sex males and freely cycling females were used. Social contact-independent (Exp. 1) or -dependent (Exp. 2) targeted helping was compared between males and females. Further, the impact of direct visualization between Target and Observer (Exp. 3) and the differences in ultrasonic vocalizations (USV, Exp. 4) were studied during social contact-independent targeted helping. Finally, subsets of animals were sacrificed, and neural activity within regions of interest was evaluated via immunohistochemistry of the immediate early gene *c-fos* (Exp. 5). **B)** Timeline for all behavioral evaluations. One rat in a cage pair was randomly assigned to be the Target or Observer. During acquisition, Observers were placed in the dry side of the chamber and given the opportunity to release the Target from the water chamber. Roles were reversed during Reversal, where rats that were Targets became Observers and vice versa. Data were collected during three timepoints: early acquisition (EA, orange), as an indication of an initial helping response; late acquisition (LA, green) to examine the effect of habituation to aiding a familiar conspecific, and reversal (Rev, blue) to see if previous experience plays an equal role in helping behavior.

## RESULTS

### Exp 1. Targeted Helping in Male and Female Rats

Groups of male and female Observers (n=21 pairs/sex) performed our lab’s social contact-independent targeted helping task to discern if any sex differences in chain pull latency were present during acquisition or reversal (**Figure 2A** depicts the timeline). For the acquisition phase, a mixed effects 2-way ANOVA was used to account for missing trials in a small subset of animals. The analysis showed a main effect of time [**Figure 2B, F** (9,344)=13.00, *p*<0.0001] and sex [F (1,40)= 7.272), *p* = 0.0102], with males having faster chain pull latencies to release their partner compared to females. However, the time x sex interaction was not significant. Post hoc analysis on the main effect of time revealed the latencies on days 2-10 were significantly faster (*p*<0.0005) compared to day 1 (**Figure 2B**). In order to determine if sex differences were present during early acquisition (EA) or late acquisition (LA), averages were taken for latencies on days 1-2 (EA) and 9-10 (LA) for each sex and subsequently compared (**Figures 2C and 2D**). Unpaired t-tests showed no significant difference between males and females at the EA or LA timepoints. Therefore, we calculated an acquisition index 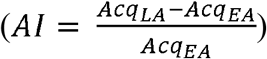for both sexes and found that the main effect of sex across acquisition was driven by a subset of low responding females (see **Supplemental Figure 1** for additional analysis).

**Figure 2.**
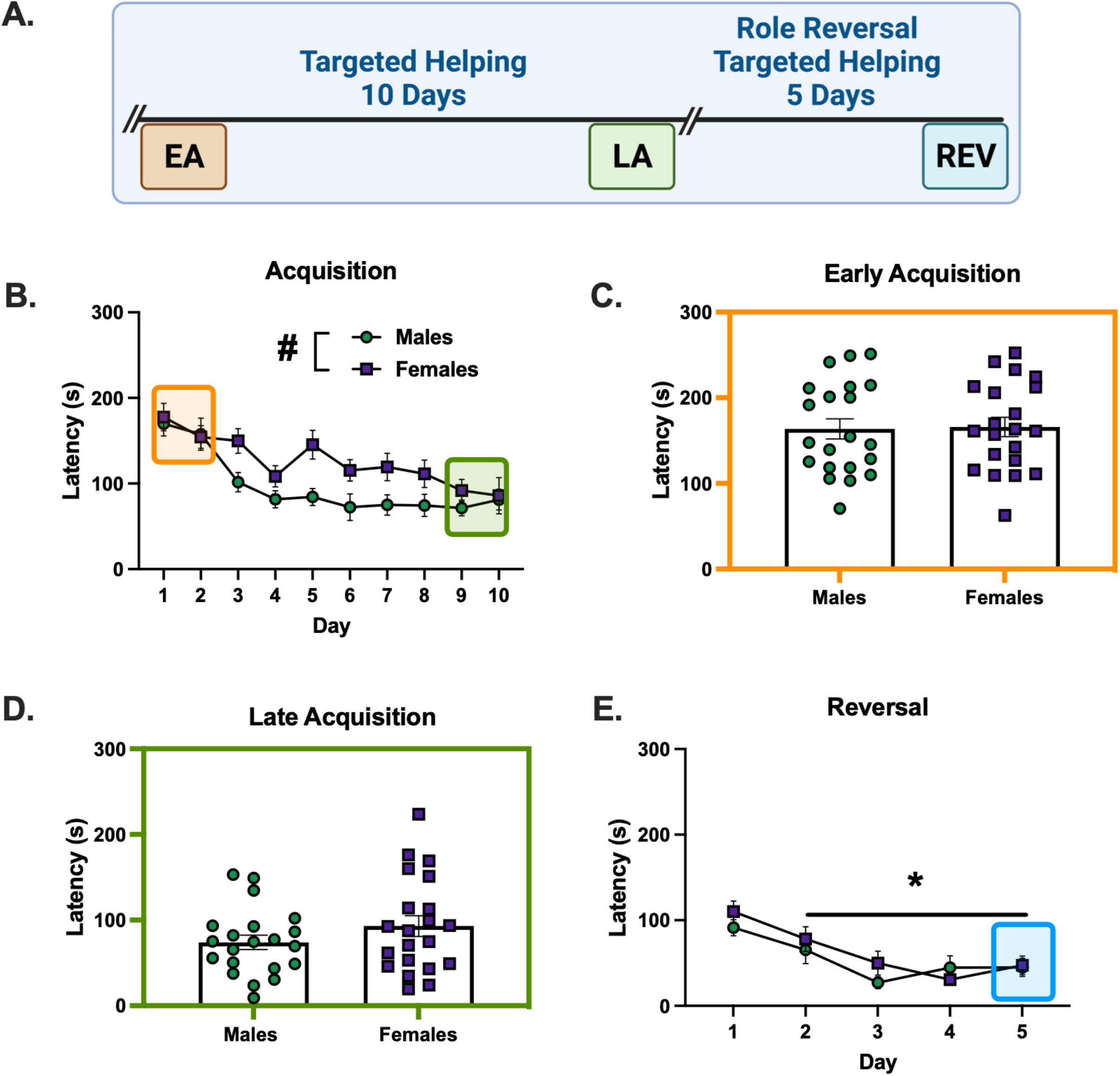
Elucidation of Sex Differences During Targeted Helping. **A)**. Performance of male and female (n=21 pairs/sex) rats during the social contact-independent targeted helping task revealed the Observers’ latency to release a distressed partner decreased over 10 days. Significantly shorter latencies occurred on days 3-10 compared to day 1. **B)**. A main effect of sex was also seen in acquisition, with male latencies being faster compared to females. **C-D)** Unpaired t-tests comparing chain pull latencies during early (EA) (**C**) and late (LA) (**D**) acquisition did not show a difference between males and females. Sex differences were also not present during task Reversal. While chain pull latency decreased, with days 2-5 significantly shorter than day 1, no main effect of sex was seen in Reversal. *Significant difference from day 1, p<0.05. #Significant difference between males and females, p<0.05.

In the Reversal (Rev) phase, there was again a main effect of time [F (4,88)=13.72, *p*<0.0001], with the latencies on days 2-5 found to be significantly faster than day 1 (p<0.05) shown in post hoc analysis of the main effect of time (**Figure 2D**). However, no effect of sex was seen between male and female R-Observers.

### Exp 2. Males and Females Readily Release a Conspecific if Social Interaction is Possible

**Figure 3A** depicts the timeline for males and females performing a social interaction task. In this task, a chain pull response released the Target into the same chamber as the Observer^42^. **Figure 3B** demonstrates that, during acquisition, males and females (n=4/sex) release a distressed conspecific in a targeted helping apparatus that affords social contact at similar rates (main effect of time [F (9, 54) =10.03, *p*<0.0001]). Post hoc analysis of the main effect showed chain pull latency on days 4-10 was significantly faster compared to day 1 (*p*<0.05). The 2-way ANOVA did not reveal a main effect of sex during acquisition. Indeed, no differences were seen in the unpaired t-tests comparing male and female latencies during EA (**Figure 3C**) or LA (**Figure 3D**). Similarly, during Rev, only a main effect of time [F (4,24) = 29.34, *p*<0.0001] was found. Specifically, Reversal days 2-5 were significantly faster compared to day 1 (*p*<0.0001) (**Figure 3E**).

**Figure 3.**
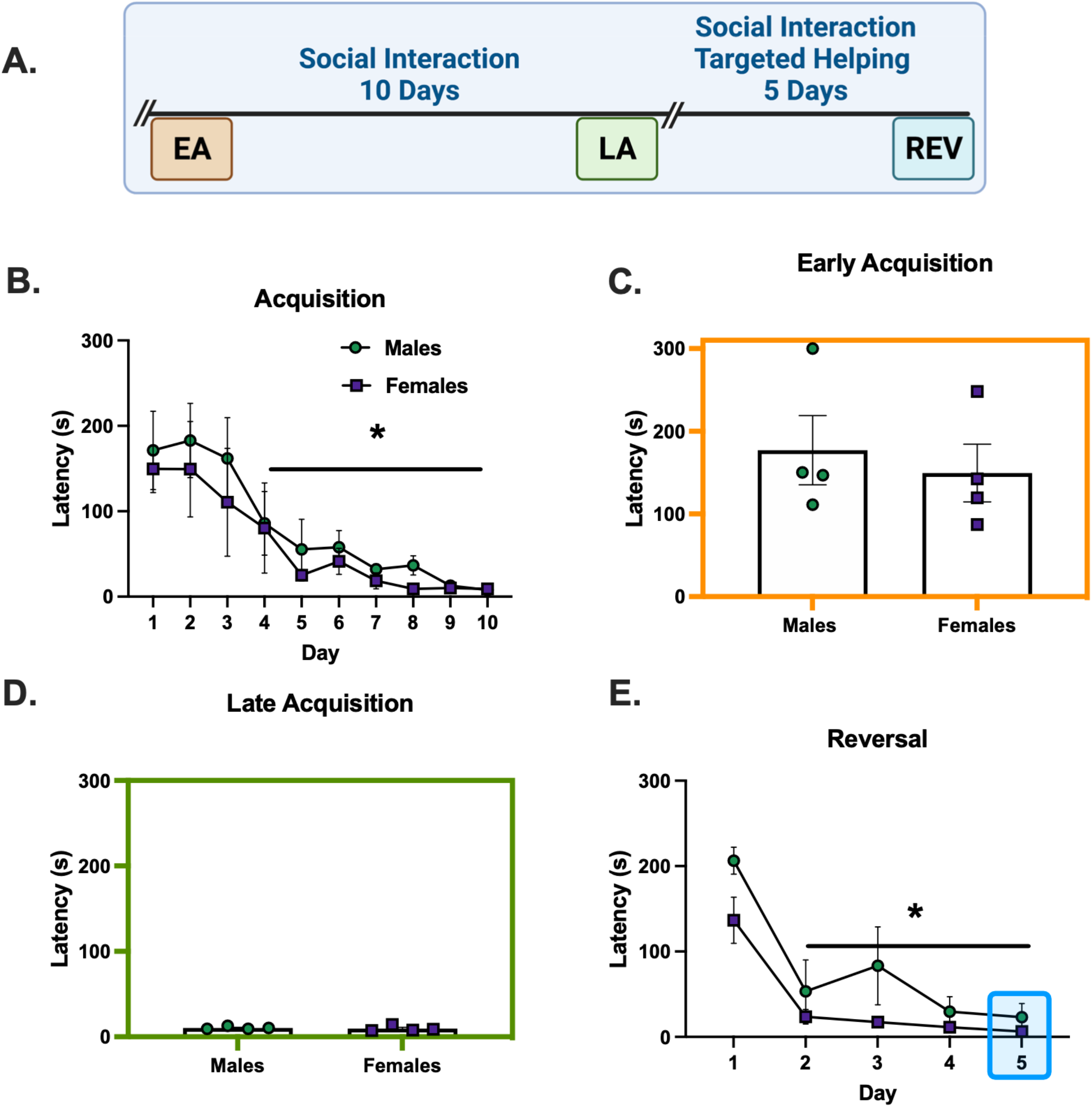
Males and Females Readily Release a Conspecific if Social Interaction is Possible. **A**) Performance of male and female (n=4 pairs/sex) rats during the helping task where social contact is possible. **B)** Chain pull latencies for males and females during acquisition did not differ; latencies decreased over time, with days 4-10 significantly faster compared to day 1. **C-D**) Unpaired t-tests comparing chain pull latencies during early (EA) (**C**) and late (LA) (**D**) acquisition did not show a difference between males and females. **E**) Latencies again decreased in reversal, specifically on days 2-5 compared to day 1. However, no main effect of sex was identified. **\***Significant difference from day 1, p<0.05.

### Exp 3. Sex Differences in Empathic Behavior when Visualization of the Conspecific is Prevented

Figure 4. shows the chain pull latency of males and females during social contact-independent targeted helping when the Plexiglas divider between the Observer and Target was obstructed. Only a main effect of time [F (9,126) = 9.733, *p*<0.0001] was revealed by the 2-way ANOVA during acquisition, with days 4-10 significantly different from day 1. There was a strong trend for an effect of sex [F (1,14) = 4.27, *p* = 0.0589, **Figure 4A**]. Unpaired t-tests comparing males and females during EA and LA revealed females were significantly faster during EA compared to males [**Figure 4B**, t (14) = 2.36, *p* = 0.043], but not during LA (**Figure 4C**). A main effect of time [F (4,56) = 4.764, *p* = 0.0022] was found during Rev when visualization of the conspecific was prevented (**Figure 4D**), but neither a main effect of sex nor a time x sex interaction was shown to be significant.

**Figure 4.**
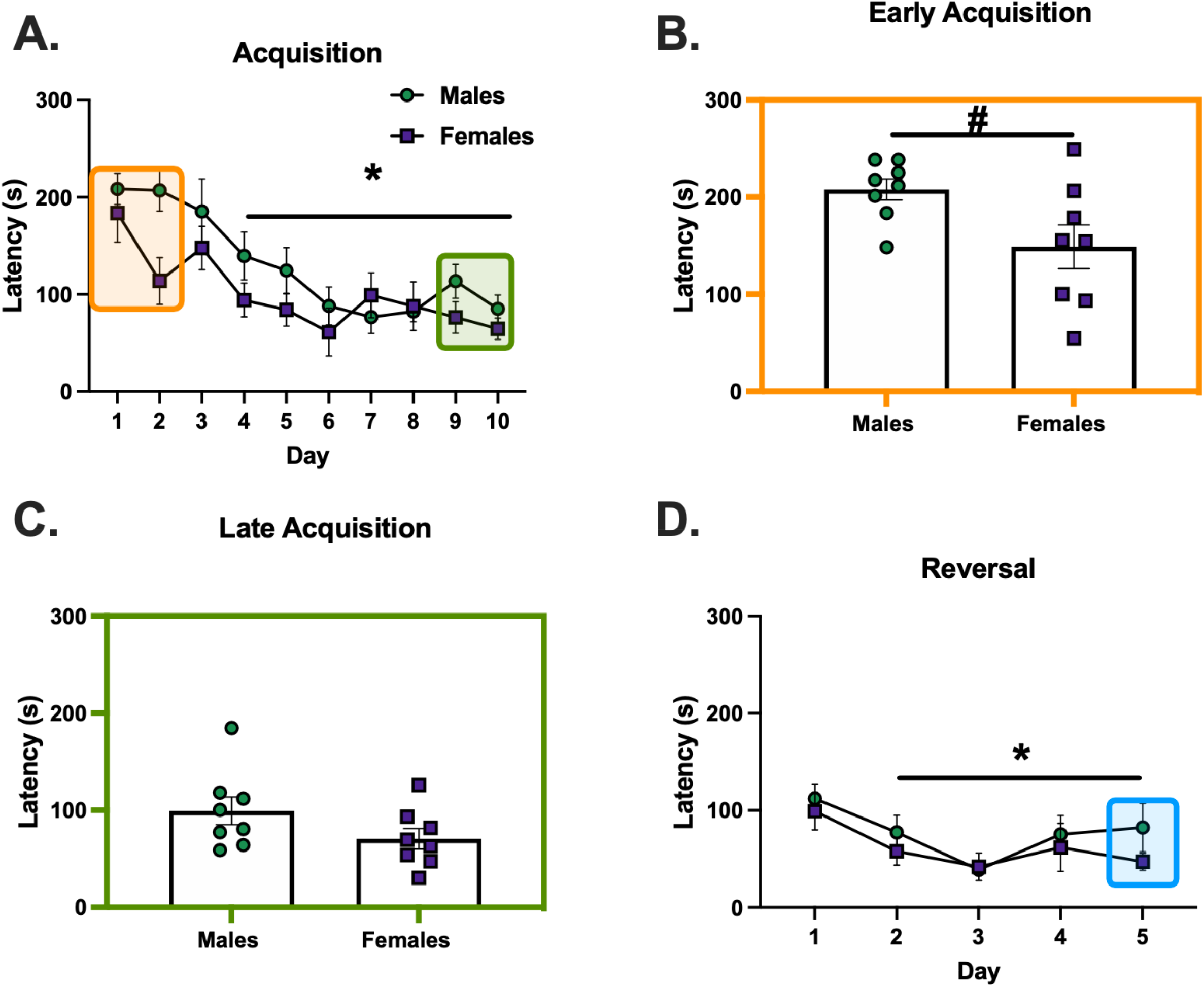
Sex Differences in Helping Behavior when Visualization of Conspecific is Prevented. **A)** Male and female chain pull latencies decreased over time, with days 4-10 significantly faster compared to day 1. There was a strong trend towards a main effect of sex, but it did not reach significance (p=0.0589). **B-C**) Unpaired t-tests were performed comparing the average latencies for early (EA) and late (LA) acquisition between males and females. **B**) Female latencies were significantly faster compared to males in EA. **C**) During LA, however, there was no sex difference in chain pull latency. **D**) While the overall latency decreased in reversal, specifically on days 2-5 compared to day 1, no main effect of sex was identified. *Significant difference from day 1, p<0.05. #Significant difference between males and females, p<0.05.

### Exp 4. Ultrasonic Vocalizations

In the next experiment, male and female rats (n=4-8 groups/sex) went through the social-independent targeted helping task (**Figure 5A**), and USV were recorded. In adult rats, two main USV have been categorized; aversive (between 18-35 kHz) calls during stressful events, and prosocial/appetitive (>35 kHz) calls^25, 45-46^, used broadly to examine rats’ affect. Samples of these calls are depicted in **Figure 5B**. To determine the range and proportion of communicative frequencies during the task in males and females, calls for each sex in EA, LA, and Rev were used to generate a frequency of distribution graph of the call frequencies (kHz). Call frequencies were binned in 5 kHz increments, and relative frequencies (% of total) were calculated for each timepoint. USV of each rat were categorized as ‘distress’ (18-35 kHz) or ‘prosocial’ (>35 kHz), and each category was analyzed as a percent of total calls using 2-way ANOVAs, with sex (male vs. female) and group (Targets vs. Observers) as the variables. During EA, the frequency distribution analysis indicated a bimodal distribution of call frequencies (**Figure 5C**). When the calls were split into ‘distress’ or ‘prosocial’, there were main effects of sex [F (1,20)=7.33, *p* =0.0136], group [F (1,20)=5.278, *p* =0.0325], and a sex x group interaction [F (1,20)=7.655, *p* =0.0119]. Post hoc analysis indicated female Targets had a significantly larger proportion of their total calls fall within the distress range, and therefore significantly fewer within the prosocial range, compared to all other groups (p<0.005) (**Figures 5D and 5E**). The same analysis was performed for males and females at the LA and Rev timepoints. There were no differences in the percent of distress or prosocial calls in male or female Observers or Targets (**Figures 5F-K**).

**Figure 5.**
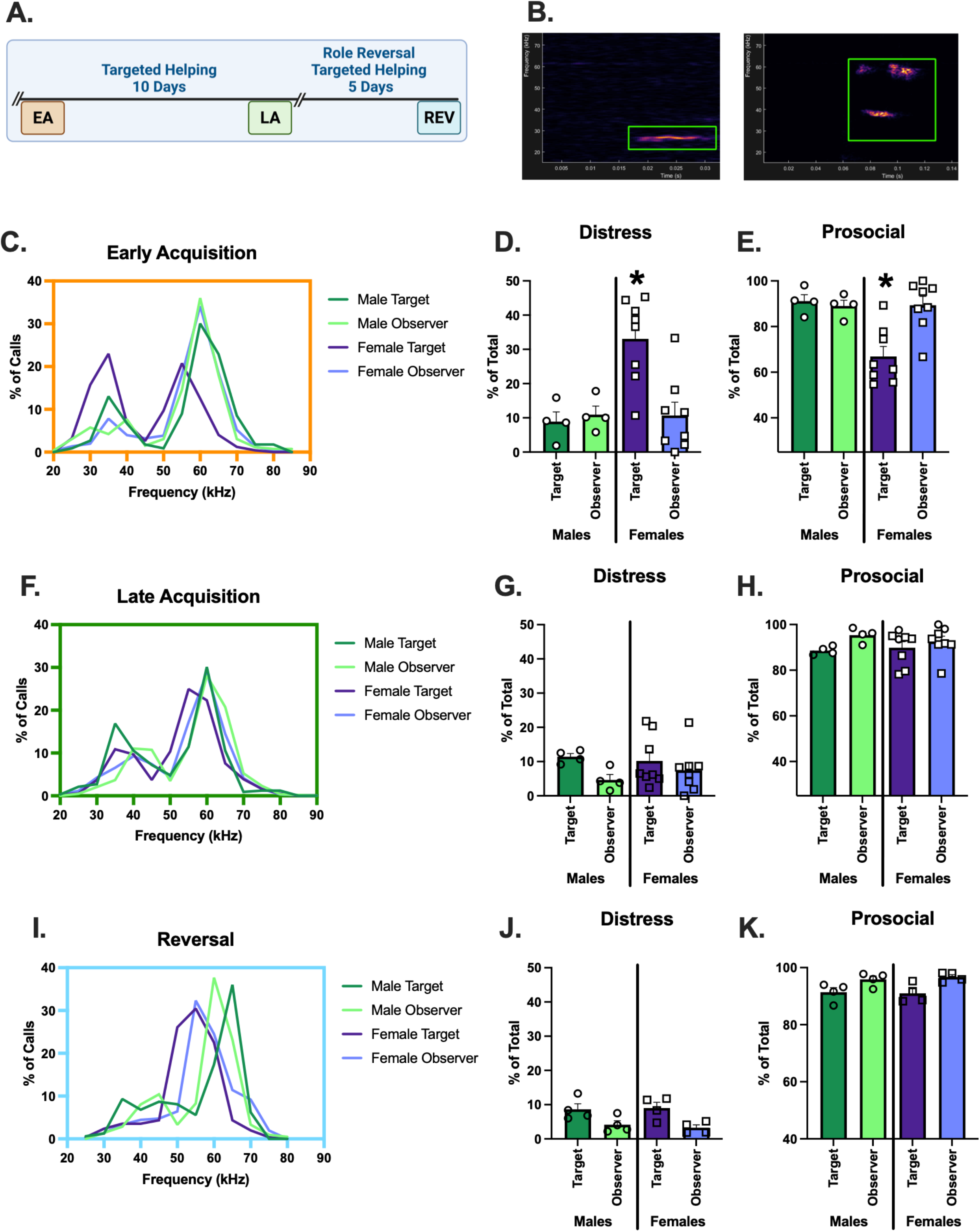
Comparison of USV Frequencies during Targeted Helping between Males and Females. **A**). Experimental timeline for helping behavior. Ultrasonic vocalizations (USV) were recorded during early (EA) and late (LA) acquisition, as well as reversal (Rev) as a proxy for the rats’ affective state during the task. USV were recorded and categorized based on frequency into two groups, ‘distress’ (18-35 kHz) or ‘prosocial’ (>35 kHz). **B**) An example of a distress and prosocial call shown in DeepSqueak software. **C-E**) USV analysis for early acquisition. **C**) Frequency of distribution graph of all calls during EA indicates a bimodal distribution of call frequencies, roughly corresponding to the ‘distress’ and ‘prosocial’ ranges, in all groups. **D-E**) An analysis of calls broken into distress or prosocial range for each group as measured by a percentage of total calls made. **D**) Female Targets make a significantly greater percentage of distress calls in EA compared to all other groups. **E**) Correspondingly, calls in the prosocial range make up a significantly smaller percentage of female Targets’ total calls compared to the other groups. **F-H**) USV analysis during late acquisition. **F**) Bimodal distribution of calls is seen in frequency of distribution graph of all calls during LA, however, fewer calls are made in the distress range. **G-H**) No differences were seen between groups in % of calls made in distress (**G**) or prosocial (**H**) ranges. **I**) There is a diminished bimodality in the frequency of distribution graph for total calls during the reversal timepoint. **J-K**) Further, no differences were seen between groups in % of calls made in distress (**J**) or prosocial (**K**) ranges.= *p<0.05

### Exp 5. C-Fos Total Count Varies Across Substrate, Group, and Sex

In order to discern the neural substrates critical during targeted helping, as well as the temporal changes and sex effects during the task, groups of male and female Observers (n=4-6) either performed the social contact-independent targeted helping task (Behaving, BEH), or remained in their homecages as controls (HCC). Importantly, HCC had the same behavioral history as the BEH rats, except they did not perform the task on test day. The behavior for these animals is described under **Methods, Exp 1**. Rats were sacrificed for Fos expression at three different timepoints; the second day of acquisition (EA), the final day of acquisition (LA), or on the final day of reversal (Rev), and the total number of Fos+ cells within each region of interest was quantified. Each timepoint was analyzed as a 3-way mixed ANOVA with sex (male, female), group (HCC, BEH), and brain region as the variables.

At EA (**Figure 6A**), there was a brain region x group interaction [F (8,96) = 19.56, p<0.0001] as well as main effects of brain region [F (8,96) = 152.6, p<0.0001] and group [F (1,13) = 69.42, p<0.0001]. There were no other main effects or interactions. Post hoc comparisons reveal that behaving animals had greater cellular activity in the AI, PL, IL, OFC, ACC, PVT, and BLA (p’s=0.047-0.0001). A heat map (**Figure 6B**) depicts this interaction by presenting the mean of each substrate during EA.

**Figure 6.**
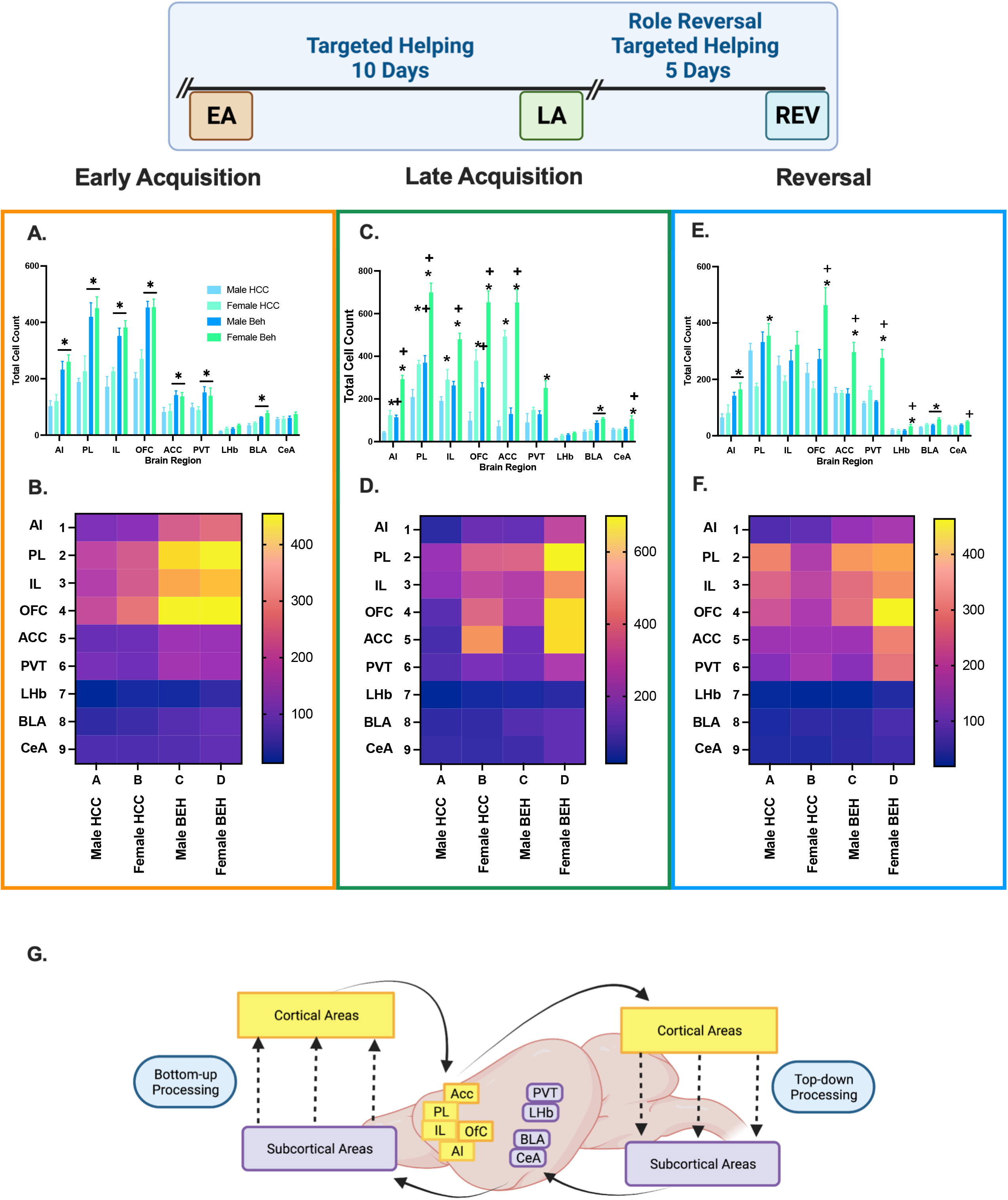
Differences in Fos+ cells across Neural Substrates of Interest. Total Fos+ cells were quantified in male and female rats that performed the targeted helping task (Behaving, BEH) or were left in their homecage on test day (HCC). Rats (4-6 groups/sex) were sacrificed at three different timepoints; the second day of acquisition (EA), the final day of acquisition (LA), or on the final day of reversal (Rev). Fos+ counts were compared in each timepoint. **A**). This figure depicts total Fos+ cell counts for male and female HCC and Beh rats during EA. **B**) Representative heat map depicting Fos+ cell means in each region of interest for each group. **C-D**) Fos activity in neural substrates during LA (**C**), with the heat map depicting mean activity in each region across the four groups (**D**). **E-F**) Fos activity in neural substrates during Rev (**E**), with the heat map depicting mean activity in each region across the four groups (**F**). *Significant difference between males and females, p<0.05 +Significant difference between BEH and HCC within the same sex, p<0.05

At LA (**Figure 6C**), there were three significant 2-way interactions: brain region x group [F (8,107) = 9.85, p<0.0001]; brain region x sex [F (8,107) = 38.15, p<0.0001]; and group x sex [F (8,16) = 8.29, p<0.0113] as well as main effects of brain region [F (8,107) = 125.1, p<0.0001], group [F (1,6) = 64.61, p<0.0001], and sex [F (1,16) = 145.8, p<0.0001]. To decompose these interactions, we conducted the analysis for each brain area separately. In the AI, there was a significant group x sex interaction [F (1,13) = 10.77, p<0.006], and a post hoc comparison shows greater AI Fos activation in females relative to males in both HCC (p<0.0075) and BEH (p<0.0001) rats. Also, greater activation was seen in BEH rats relative to HCC rats in both males (p<0.008) and females (p<0.0001). In the PL, there was a significant group x sex interaction [F (1,13) = 6.54, p<0.024] and post hoc comparisons show greater AI Fos activation in females relative to males in both HCC (p<0.018) and BEH p<0.0001) rats, as well as greater activation in BEH relative to HCC rats in both males (p<0.0.014) and females (p<0.0001). In the IL, there were main effects of group [F (1,16) = 19.71, p<0.0004] and sex [F (1,16) = 28.9, p<0.0001]. There were main effects of group [F (1,14) = 24.71, p<0.0002] and sex [F (1,14) = 61.99, p<0.0001] found in the OFC. In the ACC there were main effects of group [F (1,12) = 24.71, p<0.015] and sex [F (1,12) = 153, p<0.0001]. Main effects of group [F (1,16) = 5.81, p<0.028] and sex [F (1,16) = 9.68, p<0.007] were also found in the PVT. In the LHb there were main effects of group [F (1,15) = 11.28, p<0.0043] and sex [F (1,15) = 7.25, p<0.02]. Only a main effect of group [F (1,12) = 53.26, p<0.0001] was seen in the BLA. Finally, in the CeA, there was a significant group x sex interaction [F (1,12) = 7.23, p<0.02]. Post hoc comparison showed greater Fos activation in female BEH relative to HCC (p<0.0085) and female relative to male BEH (p<0.0001) rats. A heat map (**Figure 6D**) depicts this interaction by presenting the mean of each region during LA.

During Rev, there was a significant 3-way interaction between brain region x group x sex [F (8,103) = 3.73, p<0.0007]. To decompose these interactions, we performed the analysis of each substrate separately. In the AI, there was a significant main effect of group [F (1,13) = 15.04, p<0.0019] with BEH rats having higher activation than HCC. In the PL there was a group x sex interaction [F (1,13) = 5.07, p<0.042] and a main effect of group [F (1,13) = 9.92, p<0.0077]. Specifically, BEH females had greater Fos activity than female HCC (p<0.014). There were no differences in the IL. In the OFC there was a group x sex interaction [F (1,13) = 7.56, p=0.017] and a main effect of group [F (1,13) = 14.9, p<0.0020]. Specifically, BEH females had significantly more activation than female HCC (p<0.0021) and BEH males (p<0.033). In the OFC there was a group x sex interaction [F (1,13) = 8.99, p=0.010] and a main effect of group [F (1,13) = 8.45, p<0.0012] and sex [F (1,13) = 9.2, p<0.009]. Specifically, BEH females had significantly more neuronal activity than female HCC (p<0.0045) and BEH males (p<0.045). In the PVT there was a group x sex interaction [F (1,13) = 7.59, p=0.0164] and main effects of group [F (1,13) = 8.98, p<0.0103] and sex [F (1,13) = 26.05, p<0.002]. Specifically, BEH females had significantly more neuronal activity than female HCC (p<0.0043) and BEH males (p<0.0004). There were no differences in the lateral habenula (LHb). In the BLA there were main effects of group [F (1,12) = 15.45, p<0.0020] and sex [F (1,12) = 21.71, p<0.0006]. In the CeA there was a group x sex interaction [F (1,13) = 3.85, p=0.071] and a main effect of group [F (1,13) = 13.26, p<0.0030]. Specifically, BEH females had significantly more neuronal activity than their HCC (p<0.0079). A heat map (**Figure 6F**) depicts this interaction by presenting the mean of each region during Rev.

Noting the sex differences in HCC animals that were evident across cortical and subcortical areas, we highlight these baseline differences in **Supplemental Figure 2**. To account for relative changes in neural activation, Fos+ cell counts were adjusted to a percent of baseline and analyzed over the different timepoints (**Supplemental Figure 3**).

To link the behavior with a neurobiological endpoint, we determined the relationship between chain pull latency and neural activation across multiple brain areas during EA, LA, and Rev. In EA, a marked negative relationship emerged between Fos activation and response latency in multiple cortical areas, including: the AI (r=0.857, p<0.0054), the PL (r=0.809, p<0.0109), IL (r=0.690, p<0.0347), ACC (r=0.643, p<0.048), and OFC (r=0.649, p<0.0481). In these areas, longer response latencies were associated with reduced neural activity levels. A positive relationship also emerged in the PVT (r=0.714, p<0.0288), meaning as latencies increased, so did neural activity (**Figure 7** depicts these correlations). During LA (**Figure 8**), there were positive correlations, rather than negative, reaching significance in the ACC (r=0.690, p<0.0347) and the OFC (r=0.612, p<0.0334). There were no correlations between latency and activity of the neuronal substrates examined during the Rev trials.

**Figure 7.**
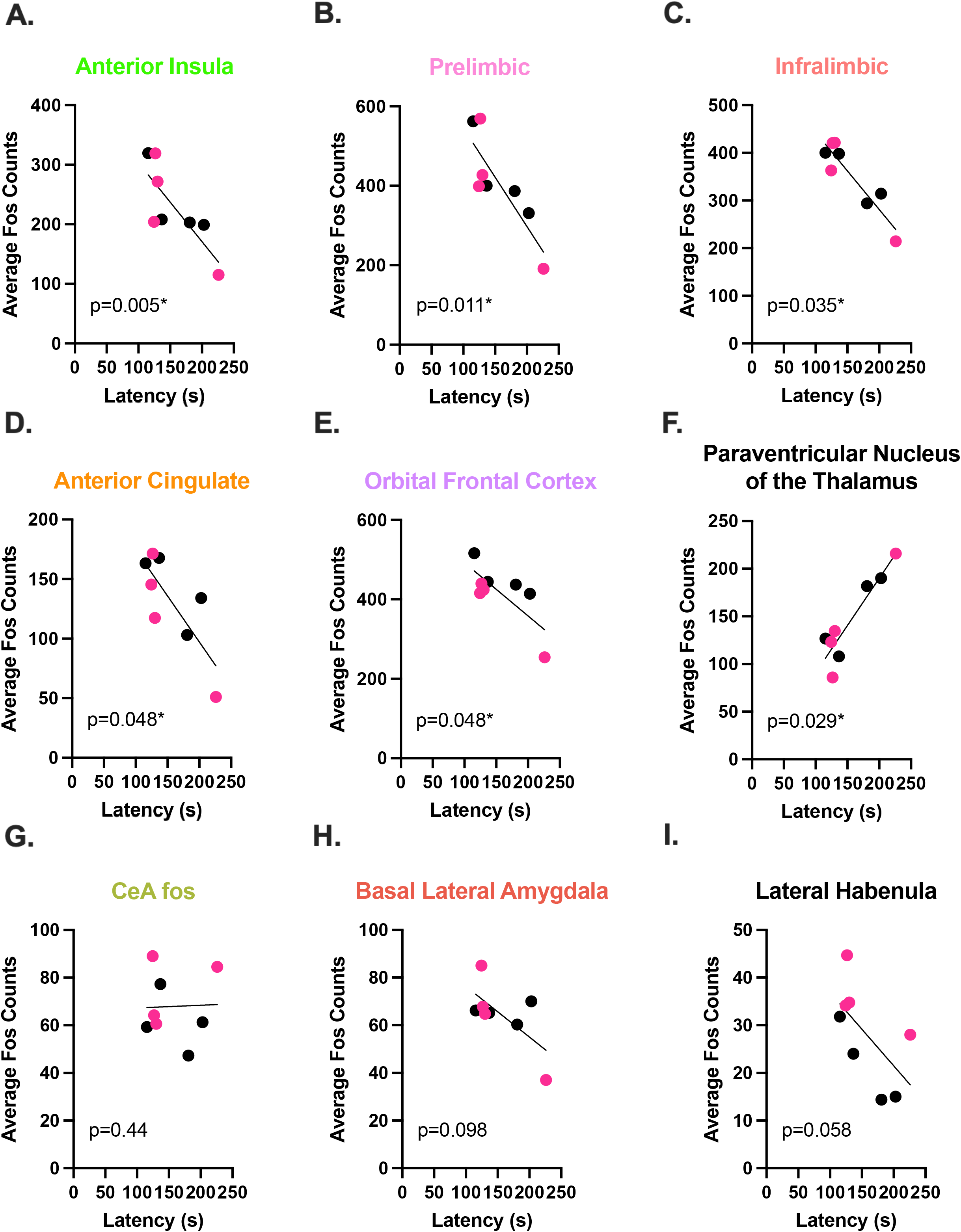
Correlation between Release Latency and Fos Activity during Early Acquisition. In order to determine a relationship between helping behavior and a neurobiological endpoint, correlational analysis was performed between the last latency and average Fos+ cell counts for male (data points in black) and female (red) Observers during early acquisition in the following regions of interest: anterior insula (**A**), prelimbic (**B**), infralimbic (**C**), anterior cingulate (**D**), and orbitofrontal (**E**) cortices, paraventricular nucleus of the thalamus (PVT) (**F**), central amygdala (**G**), basolateral amygdala (**H**), and lateral habenula (**I**). Significant negative correlations were found between final latency and mean Fos count in the five cortical regions analyzed. In contrast, a positive correlation was observed in the PVT. *Significant correlation, p<0.05

**Figure 8.**
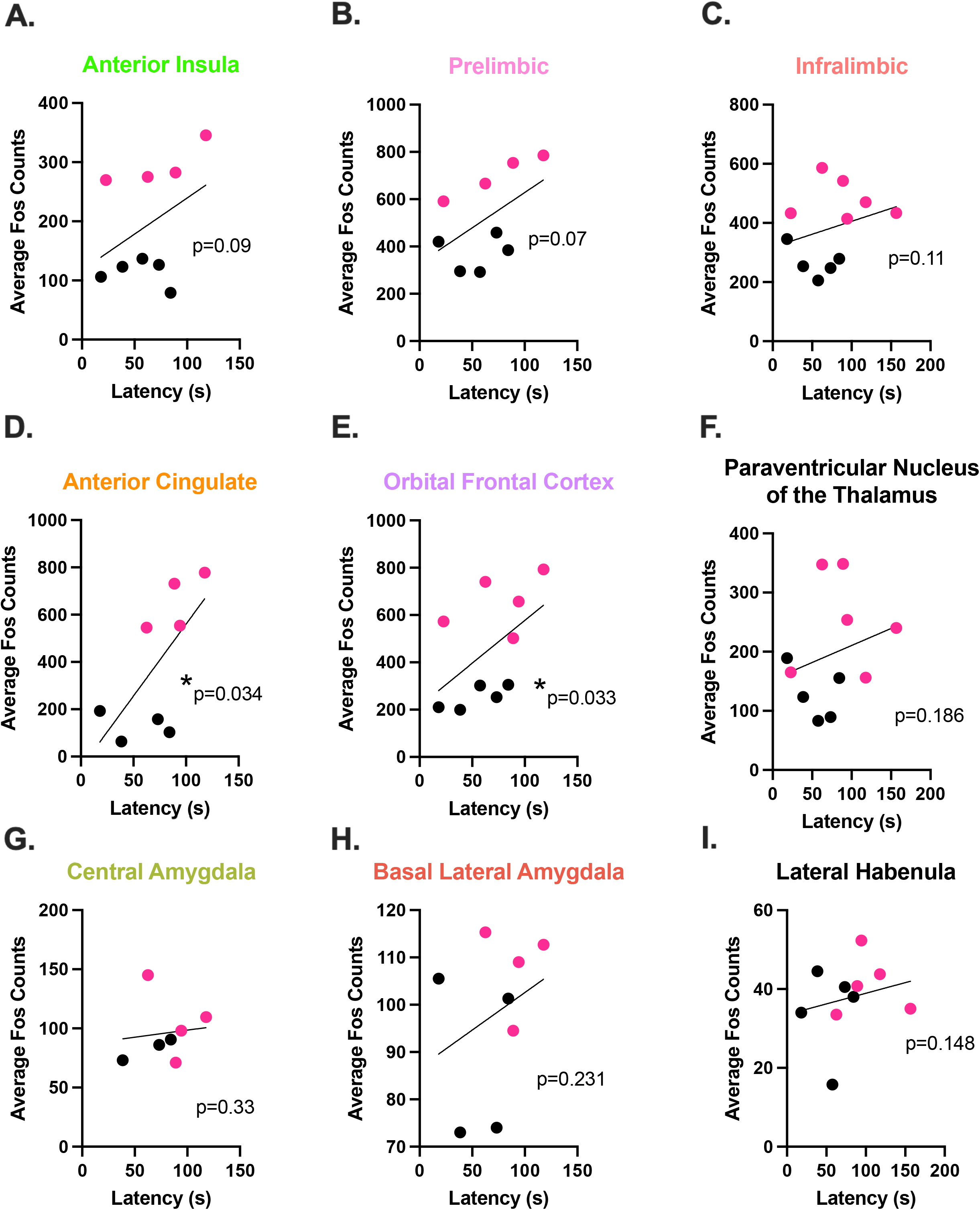
Correlation between Release Latency and Fos Activity during Late Acquisition. In order to determine a relationship between helping behavior and a neurobiological endpoint, correlational analysis was performed between the last latency and average Fos+ cell counts for male (data points in black) and female (red) Observers during late acquisition in the following regions of interest: anterior insula (**A**), prelimbic (**B**), infralimbic (**C**), anterior cingulate (**D**), and orbitofrontal (**E**) cortices, paraventricular nucleus of the thalamus (PVT) (**F**), central amygdala (**G**), basolateral amygdala (**H**), and lateral habenula (**I**). Significant positive correlations were found in the anterior cingulate and orbitofrontal cortices, with strong trends also seen in the anterior insula and prelimbic cortex. *Significant correlation, p<0.05

## CONCLUSIONS

In the current set of experiments, we sought to elucidate behavioral, sensory, affective, and neural variables of targeted helping in male and female rats. Overall, we found that males and females release a distressed conspecific at similar rates in the three timepoints evaluated, but different mechanisms or socio-emotional cues may be used to execute the response. Sex differences in behavior were present when direct visualization of the Target was blocked (**Figure 4**), as well as the expression of ultrasonic vocalizations (USV) (**Figures 5**) during early acquisition (EA). Interestingly, males and females exhibit different neuronal activation patterns during late acquisition (LA) and reversal (Rev) (**Figure 6)**, but similar patterns in EA. Across both sexes, neural activation was negatively correlated to release latency in multiple cortical areas in EA (**Figure 7**), whereas positive relationships were evident during LA (**Figure 8**), suggesting plasticity in cortical neurons over the course of targeted helping.

Canonically, clinical research has intimated females have higher levels of empathic concern compared to males^10-11^, but this broadly held assumption is now up for debate^12,47^. The few rodent experiments on prosocial behaviors that include sex as a biological variable have conflicting conclusions; targeted helping is potentiated in females compared to males in some studies^7^ while other studies using emotional contagion report no difference^13-14^. In our social contact-independent model of targeted helping, we found females to have longer release latencies relative to males when assessing acquisition in its totality (**Figure 2**). This sex difference is not attributable to a difference in the motivation to receive social contact as a reward, as latencies were similar between sexes in the model of targeted helping that affords social interaction (**Figure 3**)^42,48^. The latency difference, in fact, did not occur during the two acquisition timepoints assessed, EA and LA. To further investigate this, we used an ‘acquisition index’(AI) to calculate the change in chain pull latency over the course of acquisition (see **Supplemental Figure 1** for additional information). A more positive AI indicated minimal change in release latency over time (labeled “Low AI”), while a more negative AI score is indicative of a larger latency reduction throughout acquisition (“High AI”). Low AI males reduced chain pull latency across acquisition, whereas low AI females did not (**Supplemental Figure 1D**). Our lab and others^48-49^ use change in latency over time as a proxy for learning and mastery of a behavioral task. While it cannot be ruled out that low AI females showed reduced helping behavior compared to males, it seems less likely since no differences were seen in any of the specific timepoints evaluated. Instead, we suggest similar levels of helping behavior between the sexes. However, a subset of females may have a diminished ability to understand the association between chain pull and target’s contingencies, possibly due to differences in anxiety-like behaviors^50^, which have been suggested to play a role in targeted helping^51-53^. Additionally, the interaction of reproductive hormones over the estrous cycle has not been established in targeted helping models. One study evaluating emotional contagion as a proxy of empathic concern showed females in the diestral phase of the estrous cycle behaved similarly to males but were significantly less responsive during the estrus phase^54^. These results support the notion that circulating estrogens are correlated with reduced anxiety^55^ and warrant further study on the impact of female estrous cycle on targeted helping.

As mentioned previously, evidence indicates complex, situational, and multimodal sensory communication is required to necessitate affective transfer and targeted helping^19-20^. Females released the Target faster than males when direct visualization was obstructed, specifically in EA (**Figure 4A & 4B**). We suggest females maintain similar latencies even if sight is obstructed, while males may rely more heavily on visualization of the Target early in acquisition. In addition, during EA, female Targets emit a higher proportion of distress USV compared to male Targets (**Figure 5**). These USV data expand on the findings in the literature that show Targets made more stress calls early in acquisition when release was rare^7^. We may conclude from these combined sensory data that female Targets find the water more distressing initially compared to males. Their enhanced distress calls may be aversive to the Observers and drive shared affect between females to allow them to maintain release behaviors, while males may be more dependent on sight. In order to answer this question, future studies could use a black Plexiglas divider without a Target and instead play back distress calls to determine if it is sufficient to produce release behavior. Finally, these data also imply distress calls alone are not sufficient for the initiation and maintenance of empathic behavior^7^. The high frequency calls shown to communicate prosocial information broadly^23-24, 46^ may also drive targeted helping, even in the absence of direct social contact. Overall, these experiments indicate sex differences are present in sensory and affective communication, especially early in acquisition when release is less frequent.

Distinctive sex-specific activity patterns emerged in our analysis of Observer neural activity. During EA, male and female behaving (BEH) rats had potentiated activity, as measured by the immediate early gene *c-fos*, throughout all of the cortical areas studied, as well as in the PVT and BLA (**Figure 6**). Increased cortical activity remained stable in females through LA and Rev. Subcortical areas also had different patterns of expression. Activity in the BLA was significantly potentiated compared to HCC in both male and female BEH rats at similar rates in all three timepoints evaluated (**Figure 6**). The BLA encodes emotionally salient memories, particularly of fear^56^. In contrast, male and female rats exhibited the same pattern of activity in the CeA during EA, but during LA and Rev, female behaving rats had potentiated CeA neuronal activity compared to female HCC and their male counterparts. Amygdala activity in observer rats generally mirrors that of stressed demonstrators^19^, suggesting that the CeA is highly sensitive to the distress of others^8^.

An interesting trend emerged across both sexes when neuronal activity was correlated with latency. During EA, significant negative correlations were found between release latency and activity in all cortical areas evaluated, including the AI, PL, IL, ACC, and OFC (**Figure 7A-D**). However, the PVN of the thalamus was positively correlated with latency (**Figure 7F**). In contrast, positive correlations were seen in cortical regions, particularly the ACC and PL, during LA (**Figure 8D-E**). It has been hypothesized that empathy is comprised broadly of two main neurological processes. The first, emotionally driven “bottom-up” subcortical regions, such as the thalamus, primarily drive processes that initiate and propagate feelings of shared emotions^57^. In contrast, “top-down” cortical circuitry is capable of receiving and regulating the primary emotional information in order to generate appropriate behavioral outputs^3, 38^. While both processes are necessary for empathy, the temporal role each plays in targeted helping is still unclear. Our data suggest subcortical, specifically thalamic, activity may drive initial and relatively unregulated emotional contagion response in early acquisition, correlated to slower release latencies (**Figure 7**). With progressive trials, higher order cortical regions were shown to be directly related to the significantly attenuated release latency (**Figure 8**), suggesting a top-down regulation of the initial emotional salience to perform a directed task^3, 38^. We may conclude from our Fos data that subcortical regions are less regulated due to lower levels of cortical activity in the initial trials, which likely affords a stronger emotional response, but poor behavioral outcomes. In contrast, cortical activity drives faster response times, possibly via the regulation of subcortical substrates like the thalamus. While both processes play a role in the observed behavior, it was interesting that plasticity within cortical regions across trials was correlated with improved release latency.

Cortical activity diverged between males and females. The total activity of cortical regions in females was significantly higher compared to males during late acquisition (**Figure 6**). This elevated Fos expression in females also occurred in HCC females relative to HCC males. When directly comparing the percent change from homecage control (HCC) animals, we saw females either retained or increased their relative cortical activity across time, while male cortical activity generally diminished over time (**Supplemental Figure 3**). As the cortical regions studied in these experiments have been shown to be necessary for cognition, decision-making, affect-sharing, emotional regulation, and perspective-taking^3, 33^-^34, 58^, it is possible females find the distress of the conspecific more salient and necessitates potentiated cortical activity compared to males for the same behavioral outcome.

A recent study by Ben-Ami Bartal and colleagues^41^ extensively explored the neural response of rats during prosocial release task. They also identified similar cortical areas were potentiated during the task compared to baseline levels. A unique aspect of their study focused on neural substrates that varied depending on prosocial intent. For example, the AI, ventral and lateral OFC, and ACC were all potentiated in rats that released both ingroup and outgroup conspecifics, indicating these regions respond to a distressed rat regardless of social context^41^. In contrast, the PL and medial OFC were potentiated during targeted helping of ingroup compared to outgroup animals and baseline. The authors concluded that these regions played a specific role in the prosocial response towards ingroup animals and not to a trapped animal or social exposure alone^41^. In our studies, the distressed Targets were always the Observers’ cagemates, meaning we did not evaluate for differences between social group. However, our results seem to corroborate the importance of these cortical substrates in the prosocial aspect of targeted helping.

In conclusion, this set of experiments sought to elucidate the role of sex on targeted helping, along with sensory, affective, and neural components that may contribute to helping behavior. We believe that a convergent sex effect may be present in targeted helping, in which males and females exhibit similar behavioral outcomes, but multiple and disparate variables work together to generate the behavior^59^. Additionally, empathic sex differences previously understood to be canon are likely dependent on contextual cues and task type^60^. Subtle nuances, like housing^61^, estrous cycle^55^, and chronicity of the task must be considered when any affective or physiological variable is being studied as it pertains to helping behaviors. Overall, we believe this study lays a groundwork for future studies to explore unique social and biological variables that drive empathic behavior in both males and females.

## MATERIALS AND METHODS

### Animals

Size-matched male and female Sprague Dawley rats (n=33 pairs/sex) weighing 250-275g were pair-housed with the same sex on a 12-hour reversed light cycle (lights on at 1800). The breakdown of animals used is visualized in **Figure 1**. Animals were given food and water *ad libitum* until behavioral testing when they were then switched to a daily stable intake (20g) of rat chow (Harlan). Rats were given at least 5 days to acclimate to their cagemate. Following acclimation, one rat was randomly selected to be the “Observer” and the other the “Target.” Animals were handled and weighed for 2 days, 5 min/day before the behavioral assessment. For all behavioral evaluations, rats were transported to the experiment room and left undisturbed for 5 minutes. The tasks were performed in a sound-attenuated room with the lights off except for a single lamp used for the experimenter to view the test. All experimental procedures were conducted in accordance with the “Guide for the Care and Use of Laboratory Rats” (Institute of Laboratory Animal Resources on Life Sciences, National Research Council) and approved by The Institutional Animal Care and Use Committee (IACUC) of the Medical University of South Carolina.

### Behavioral Testing

#### Social Contact-Dependent Targeted Helping Task

Evaluation of targeted helping with the opportunity for social reward was performed in males and females (**Exp. 2**, n=4 pairs/sex) in a custom-made operant box (34.2×33.9×30.5 cm) by Med Associates (Fairfax, VT, USA). Targets were placed in 100 mm of water in the wet compartment of the apparatus, while the Observer was placed on a dry platform with access to a chain that, when pulled, opened an automatic guillotine door. Door opening allowed Targets to be released into the same dry compartment as the Observer^42^.

#### Social Contact-Independent Targeted Helping Task

In the remainder of the experiments, social interaction-independent helping behavior was evaluated (n=21pairs/sex) using a custom (Med Associates; Fairfax, VT, USA) operant box developed with three chambers^42^, as seen in **Figure 1A**. In this apparatus, Targets were placed in 100 mm of water in the wet compartment and Observers on a dry platform with access to a chain that opened an automated door. The Target was released into a dry compartment separate from the Observer. In order to determine the importance of the Observer visualizing the Target to learn to release the distressed conspecific, a separate cohort of male and female rats (**Exp. 3**, n=8 pairs/sex) were tested in the same operant box where the Plexiglas divider present between Observer and Target was painted black to prevent either animal from seeing through it.

In all experiments, latency to chain pull was taken as an index of helping behavior. Trials (20 total across 10 days, labeled “Acquisition”) lasted a total of 300 s (5 min) regardless of the chain pull latency. If the Observer did not pull the chain within the allotted time, the experimenter ended the trial and released the Target. In some of the following experiments, the role of each rat in a pair was subsequently reversed (“Reversal” phase), such that a Target became the Observer (labeled “R-observer”), and the Observer becomes the Target (“R-target”). This reversal phase was carried out for 5 days (10 trials) to determine the importance of previous experience on release latency. Two trials were conducted daily during the rats’ dark cycle^42^.

### Immunohistochemistry (IHC)

Groups of male and female Observer rats either performed the social contact-independent targeted helping task (Behaving, BEH) or remained in their homecages as controls (HCC). Importantly, HCCs had the same behavioral history as the BEH rats, except they did not perform the targeted helping task on test day. A subset of subjects was sacrificed and perfused approximately 90 minutes following the social contact-independent targeted helping task at three different timepoints (EA, LA, and Rev, n=4-6/group), and brains were collected. Briefly, rats were anesthetized with Equithesin and then transcardially perfused with 150-200 mL cold 0.9% saline followed by 400-500 mL of 10% buffered formalin. Brains were removed and postfixed in 10% formalin for 24 hours, submerged in 20% sucrose/0.1% sodium azide solution for 48 hours, and then sectioned into 50 µm tissue sections. For Fos expression, tissue sections were incubated in a rabbit anti-Fos primary antibody (Millipore; 1:1000; RRID 310107) overnight, followed by a 2-hour incubation in donkey anti-rabbit secondary antibody (Jackson ImmunoResearch; 1:500; RRID 2340584) amplified with an avidin biotin complex method (Thermo Scientific, Waltham, MA). The sections were then visualized with 3,3’ diaminobenzidine (DAB, Sigma-Aldrich, St. Louis, MO) + nickel ammonium sulfate to produce a blue-black nuclear reaction product. Slices were coverslipped using Permount, and regions of interest were photographed at 10x magnification using a Leica microscope and VideoToolbox software.

#### IHC Quantification and Analysis

For the DAB stain, blue-black nuclear immunoprecipitate from Fos-positive cells in regions of interest were quantified using a brain atlas for comparison^62^. Fos-positive (Fos+) cells that fell within each region of interest were automatically counted using a macro and averaged across sections for each rat. On average, 3 bilateral sequential sections for each region were used for analysis. Anterior-posterior coordinates for each analyzed region are as follows: PL: 3.7 to 2.7; IL: 3.2 to 2.2; ACC: 3.2 to 2.2; OFC: 4.2 to 2.7; LHb: -2.5 to -3.3; PVT: -2.5 to -3.6; BLA: -2.5 to -3.3; CeA: -2.5 to -2.8. In order to compare the change in Fos+ cells between sex and across time, each group was compared to their own homecage control (HCC) by calculating a percent change from HCC in the analysis. All images were quantified using ImageJ software (NIH).

### Ultrasonic Vocalization Detection and Analysis

Ultrasonic vocalizations (USV) were recorded in the same subset of male and female (n=4-8) rats used for behavioral and Fos analysis to understand their affective states and level of communication during targeted helping. Two high-quality condenser microphones (Avisoft Bioacoustics) were fastened to the lids of the operant box, one on the Observer’s dry side and one on the Target’s wet side. The microphones were connected to Avisoft UltraSoundGate 416Hb multichannel recording system and processed using Avisoft-SASLab Pro software (Avisoft Bioacoustics, Glienicke, Germany). USV weren recorded for one complete trial (300s) at three different timepoints (EA, LA, and Rev) with a sampling rate of 250kHz, and analyzed with DeepSqueak version 2.6.0^63^ in MATLAB. Due to background noise during the task, post-hoc denoising was carried out and subsequently rechecked for errors by an experimenter. Calls with tonality of <0.35 were considered to be background and were rejected manually from analysis. Remaining USV were reviewed by an experimenter blind to the conditions and calls that were picked up in both microphones were assigned to a particular rat by comparing power and timing of the USV across both channels.

## Data Analysis

Two-way mixed analysis of variance (ANOVAs) were used to compare the latency of chain pulls responses for social independent (**Exp. 1**) and dependent (**Exp. 2**) targeted helping as well as targeted helping when vision is obstructed (**Exp. 3)**. The between-subject variable was sex (males vs. females), with the repeated measure being days (1-10 Acquisition and 1-5 Reversal). Comparison of the timepoints defined as EA (acquisition days 1-2), LA (acquisition days 9-10), and Rev (reversal days 4-5) was performed with unpaired t-test between males and females. USV data were analyzed with two-way between subjects ANOVAs for total call counts, as well as % of total distress prosocial calls, with sex and group as the variables. Three-way mixed variable ANOVAs were used to analyze total cell counts during EA, LA, and Rev with Brain Region x Sex x Group as the independent variables. 2-way between subjects ANOVAs with sex (male vs. female) and time (EA vs. LA vs. Rev) as the independent variables were used to analyze percent change from HCC comparisons. When correlations were calculated, total Fos count was correlated with release latency at all three timepoints separately using a Spearman R correlational analysis. All post hoc comparisons were conducted using a Holm-Sidak’s correction for family wise error when appropriate, with the alpha set at 0.05. Mixed effect models were used when necessary to account for any missing data points. All analyses were conducted with Prism Software version 9.0. Unless noted, all data are expressed as the mean ± SEM.

## Acknowledgments

The authors would like to thank Jordan Carter and Jordan Hopkins for their assistance in acquiring preliminary data for the completion of this manuscript. We would also like to thank Dr. Kevin Coffey for his assistance with the DeepSqueak program.

## Author Contributions

SSC and CMR designed and ran experiments with the assistance of BJB, SJB, and AMK. Data and image acquisition was performed by SSC, BJB, and SKW, and analysis was performed by SSC with the assistance of BJB and SKW. The manuscript was written by SSC and CMR and was corrected by BJB, SJB, AMK, and CMR.

## Data Availability

The datasets generated and/or analyzed in the studies described in this manuscript are not publicly available due the ongoing nature of components of this project, but are available from the corresponding author upon reasonable request.

