## Supplemental figures 1-4 for "Neuronal, Affective, and Sensory Correlates of Targeted Helping Behavior in Male and Female Sprague Dawley Rats"

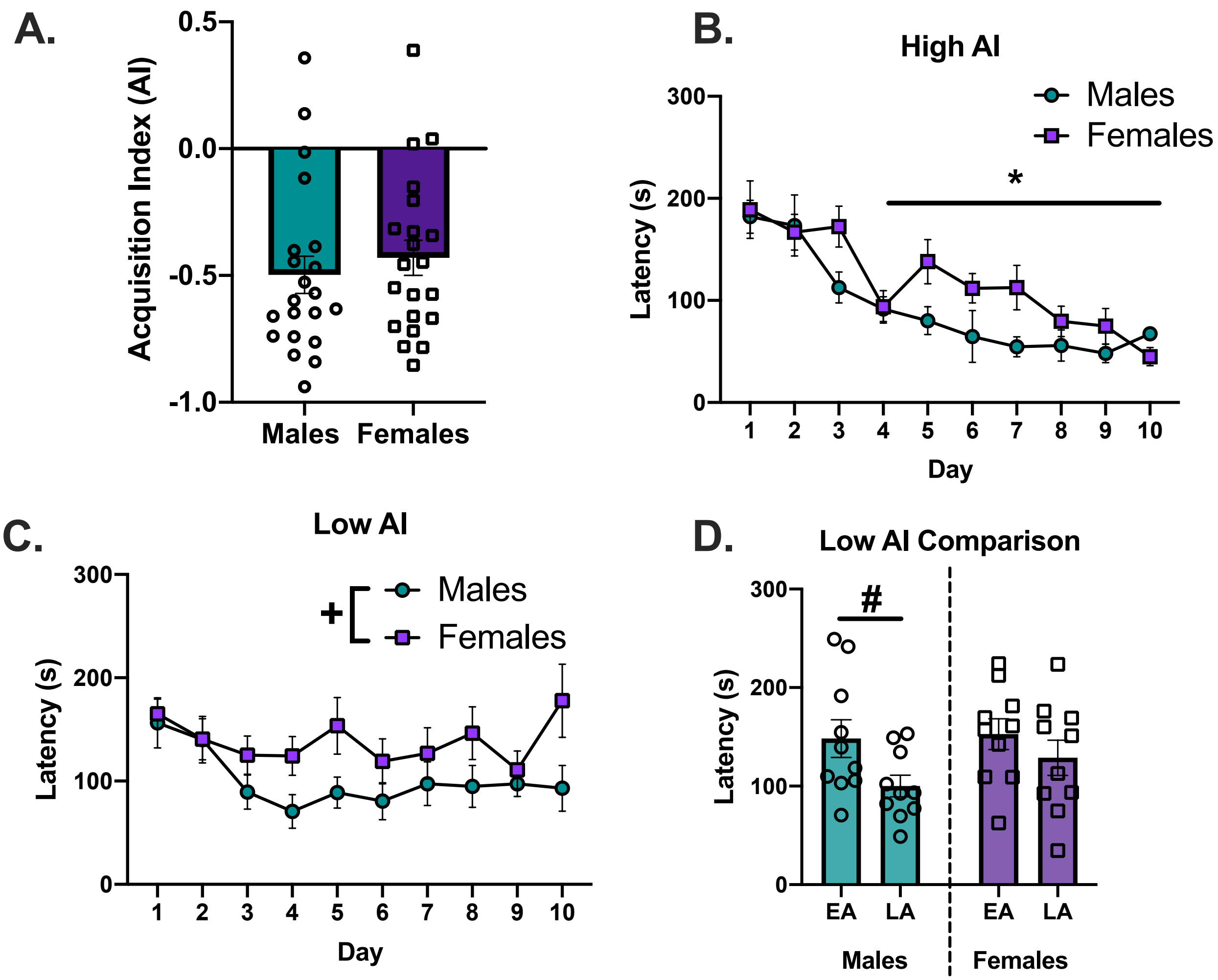


**Supplemental Figure 1: Comparison of Acquisition Index Calculation Between Sex**. In order to better understand individual and sex differences in targeted helping as measured by chain pull latency, an Acquisition Index was calculated $AI = \frac{Acq_{LA} - Acq_{EA}}{Acq_{EA}}$, where $Acq_{EA}$ is an average latency of the first 2 days of acquisition used as baseline, and $Acq_{LA}$ is an average of the final 2 days of acquisition. Positive AI reflects a reduction in latency over the course of acquisition, with AI=1 the maximum possible given each animal's baseline. **A)** An unpaired t-test comparing the AI of all males compared to all females revealed no significant difference. **B-C)** The median of each sex's AI was then used to divide each sex into two groups, "Low AI" and "High AI"[64]. A mixed effects 3-way ANOVA comparing all of these groups indicated main effects of time [$F$ (9,173) = 16.47, $p$<0.0001], sex [$F$ (1,197) = 24.61, $p$<0.0001], and index group [$F$ (1,167) = 5.091, $p$ = 0.0253]. Further, a significant time x index group [$F$ (9,167) = 3.935, $p$ = 0.0001] interaction was seen. Low and high AI groups of each sex were then compared separately. **B)** A mixed effects 2-way ANOVA comparing high AI male and female rats only showed a main effect of time [$F$ (9,173) = 16.47, $p$<0.0001], with days 4-10 significantly faster compared to day 1. **C)** The same analysis of males and females with low AI showed a main effect of time [$F$ (9,153) = 2.692, p = 0.0062] and sex [$F$ (1,18) = 5.23, $p$ = 0.0345]; however, only days 4 and 6 were significantly different from day 1. **D)** To further explore the sex effect within the Low AI group, a 2-way ANOVA was run to compare the change in latency between early (EA) and late acquisition (LA) of Low AI animals of each sex. A main effect of time was discovered [F (1,18) = 10.01, $p$ = 0.0054], and a post hoc of this effect revealed that chain pull latency decreased significantly only from EA to LA in males (p=0.0159). No difference in latency was seen in females. Taken together, these data indicate the main effect of sex in acquisition seems to be driven primarily by Low AI females that do not seem to acquire the task, as measured by a change in latency over time.

*Significant difference from day 1, p<0.05.
#Significant difference between EA and LA, p<0.05
+Significant difference between males and females, p<0.05

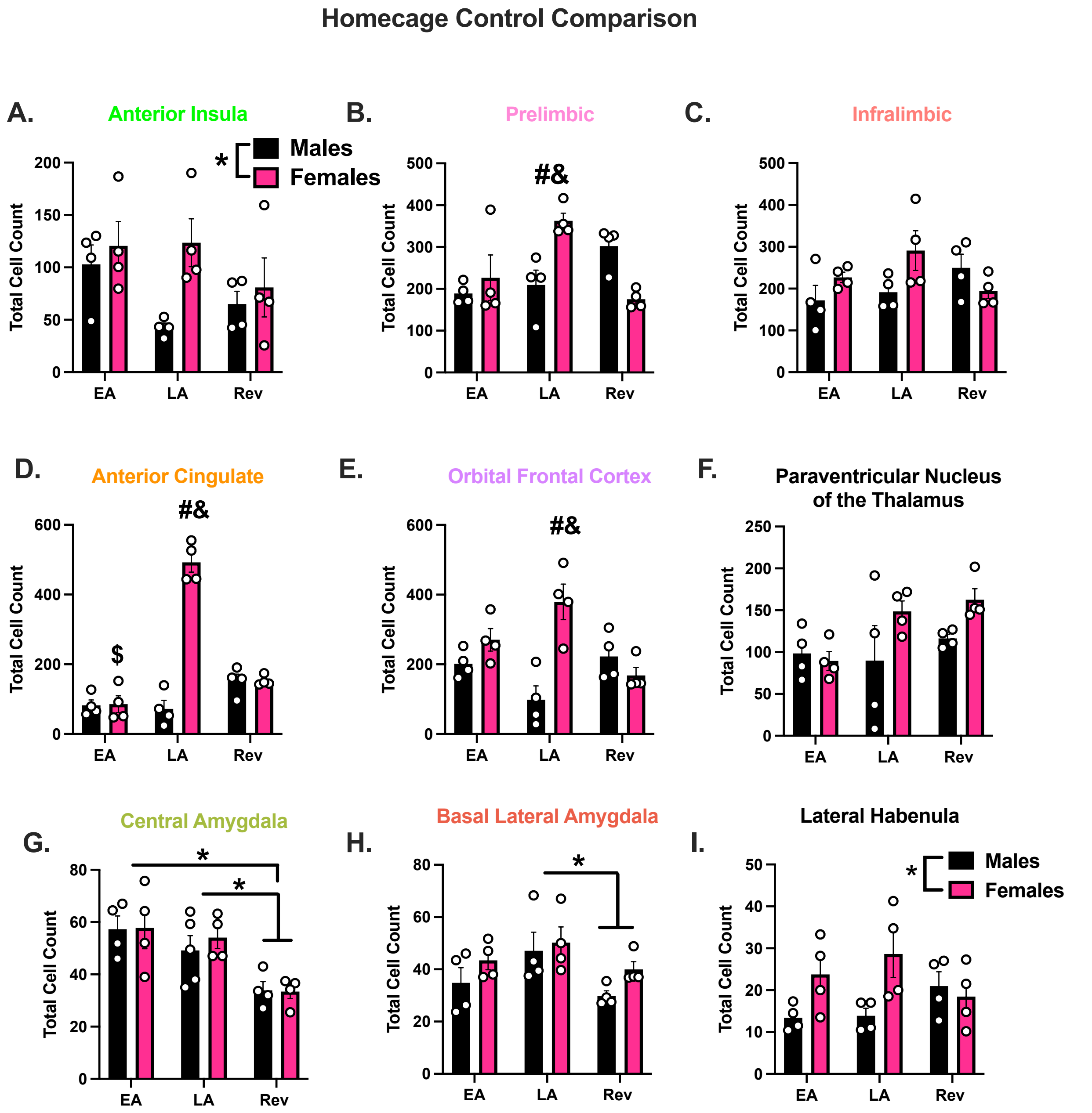


**Supplemental Figure 2**. **Sex Differences in Homecage Controls within Neural Substrates of Interest.** Groups of male and female rats (n=4/sex) acting as homecage controls (HCC) were sacrificed at one of three-timepoints; early acquisition (EA), late acquisition (LA), or reversal (Rev). Total Fos+ cells for HCC rats were counted in each region of interest and compared at each timepoint. A 2-way ANOVA was calculated, with the variables being sex (males vs. females) and time (EA vs. LA. vs. Rev). Post hocs that were reported specifically focused on comparing across sex at the same timepoint and within sex across all timepoints. **A**) The 2-way ANOVA in the insular cortex revealed a significant effect of sex [F (1,18) = 5.53, $p$ = 0.0303], but neither a main effect of time nor an interaction was found. **B**) Within the prelimbic cortex (PL), the analysis revealed a significant sex x time interaction [ F (2,18) = 10.83, $p$ = 0.0008]. Post hoc comparisons of note showed differences in total Fos+ cells between LA males and females ($p$ = 0.0275) and LA and Rev females ($p$ = 0.0055). **C**) Although there was a strong trend (p = 0.0508) for a sex x time interaction in the infralimbic cortex (IL), nothing significant was seen in the analysis. **D**) Within the anterior cingulate cortex (ACC), relevant post hocs concluded Fos+ cells for females at the LA timepoint were significantly higher than males at LA, as well as females at both EA and Rev. **E**) Analysis in the orbitofrontal cortex (OFC) revealed females at LA had significantly higher Fos+ cell counts compared to LA males, as well as Rev females. **F**) Neither a main effect of sex nor a main effect of time was observed in the analysis for the paraventricular nucleus of the thalamus (PVT). **G**) Within the central amygdala (CeA), post hoc analysis of the main effect of time [$F(2,19) = 11.05$, $p = 0.0007$] revealed Fos+ cell counts at the EA timepoint were significantly higher than both LA and Rev. **H**) Similarly, within the basolateral amygdala (BLA), post hoc analysis on the main effect of time [$F(2,18) = 4.10$, $p = 0.0341$] showed Fos+ cells counts in Rev to be significantly lower compared to LA. **I**) Only a main effect of sex was seen in the lateral habenula (LHb) [$F(1,18) = 6.215$, $p = 0.0226$].

* $p<0.05$
**#**Significant difference from same timepoint of other sex, $p<0.05$
**&**Significant difference from reversal timepoint of own sex, $p<0.05$
**$**Significant difference from late acquisition of own sex, $p<0.05$

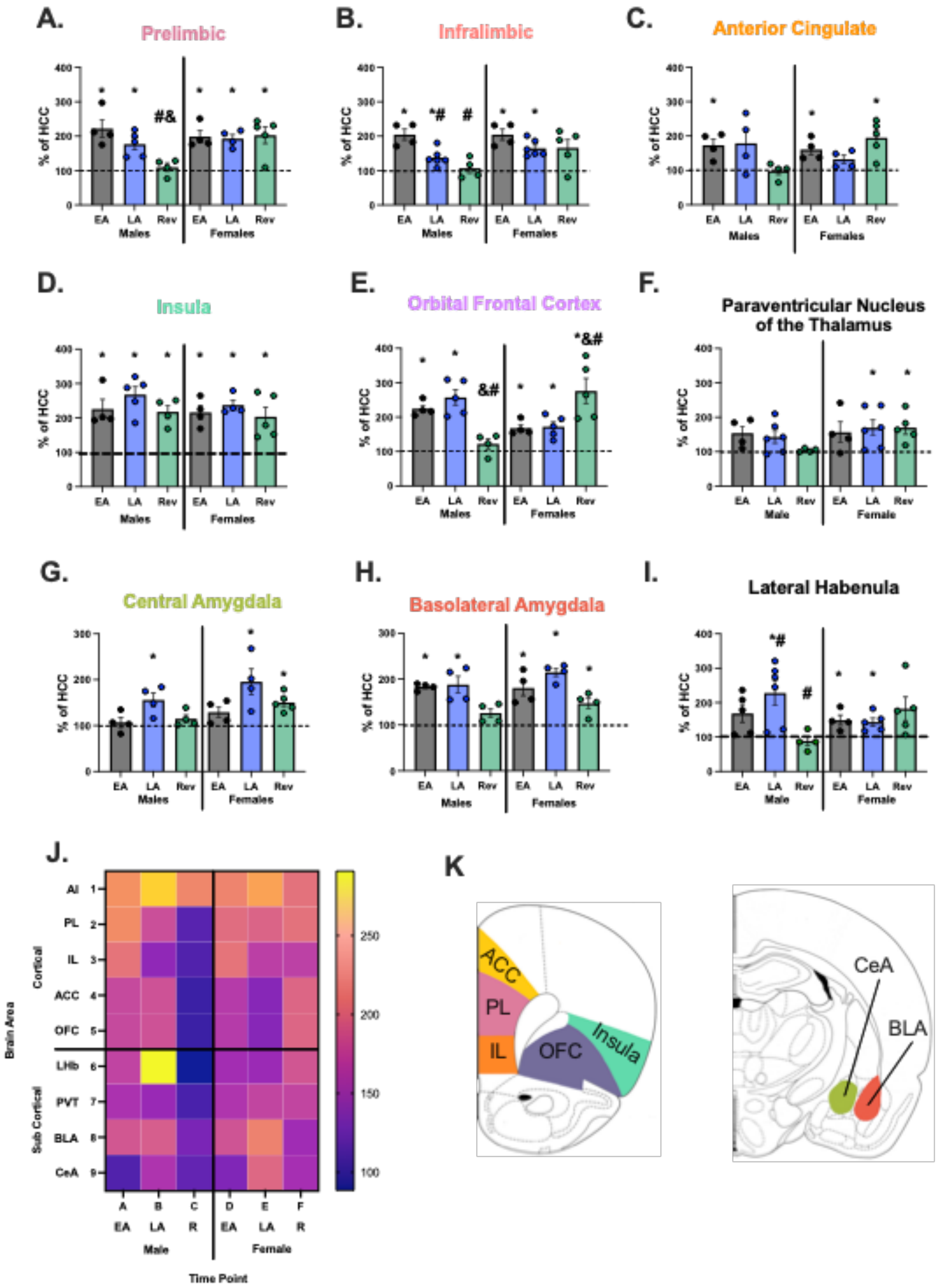

A.
Prelimbic
B.
Infralimbic
C.
Anterior Cingulate
D.
Insula
E.
Orbital Frontal Cortex
F.
Paraventricular Nucleus of the Thalamus
G.
Central Amygdala
H.
Basolateral Amygdala
I.
Lateral Habenula
% of HCC
EA LA Rev EA LA Rev
Males Females
Male Female
J.
Brain Area
Cortical
Sub Cortical
AI 1
PL 2
IL 3
ACC 4
OFC 5
LHb 6
PVT 7
BLA 8
CeA 9
A B C D E F
EA LA R EA LA R
Male Female
Time Point
K
ACC
PL
IL
OFC
Insula
CeA
BLA

**Supplemental Figure 3. Percent Change Differences in Fos+ cells across Neural Regions of Interest.** Male and female rats seem to exhibit a similar pattern of behavior during the task during EA, LA, and Rev (**Figure 2**), but there is a sex difference inherent in the baseline activity of homecage control (HCC) animals between sex in many of the regions examined (**Supplemental Figure 2**), suggesting a quantitative sex effect. This effect may be producing the convergent effect seen in behavior, in which males and females exhibit similar targeted helping, but the underlying process mediating it differs[59]. Therefore, to better compare Fos levels between sexes, we converted the total Fos+ cell counts into a percent of HCC for each group using the following formula: $\%HCC = \frac{Fos_{BEH}}{Fos_{HCC}}$, with Fos+$_{EMP}$ being the total Fos+ cell count in the BEH group, and Fos+$_{HCC}$ the total in the respective HCC animals. Values are set to 100% of HCC. That way, the overall sex differences in basal neural activity can be accounted for, and the sexes can be directly compared between the three timepoints. For each neural substrate of interest, a 2-way ANOVA was performed, and where appropriate, post hocs comparing each timepoint (EA vs. LA vs. Rev) were compared within sex to understand if the trend in activity across time in each region was different in males compared to females. **A**) Within the prelimbic cortex, a main effect of time [F (2,20) = 3.839, *p* = 0.039], as well as a significant interaction [F (2,20) = 4.605, *p* = 0.023] was found. Post hoc analysis revealed male Rev rats had a significantly attenuated change of Fos+ cells from HCC as compared to EA (*p* = 0.0024) or LA (*p* = 0.0483) males. In contrast, female timepoints were not significantly different from one another. **B**) The infralimbic cortex analysis showed main effects of both time [F (2,23) = 9.526, *p* = 0.001] and sex [F (1,23) = 5.127, *p* = 0.0333]. Specifically, EA males had a potentiated % change in Fos+ cells compared to both LA males (*p* = 0.0108) and Rev males (*p* = 0.0013), with no changes within females. **C**) Main effects of sex [F (1,18) = 66.75, *p*<0.0001] and time [F (2,18) = 45.27, *p*<0.0001], along with a significant group x time interaction [F (2,18) =64.79, *p*<0.0001] were seen in the anterior cingulate cortex. Pertinent post hoc analyses showed that total counts in LA females were significantly higher than those of LA males (*p*<0.0001), LA females (*p*<0.0001), and Rev females (*p*<0.0001). **D**) An approximate two-fold increase in Fos+ cells compared to HCC was seen within the insula across all groups. Therefore, nothing of significance emerged from the analysis. However, one-sample t-tests indicated that activity in both sexes at all timepoints was significantly higher (p<0.05) compared to HCC. **E**) The 2-way ANOVA of the orbitofrontal cortex (OFC) showed a main effect of sex [F (1,18) = 11.96, *p* = 0.0028] and significant sex x time interaction [F (2,18) = 11.90, *p* = 0.0005]. Specifically, LA females had a higher Fos+ count than LA males (*p* = 0.0003) and Rev females (*p* = 0.0061). **F**) Within the PVT, there was a trend towards a main effect of sex (*p* = 0.083), but nothing significant was revealed in the analysis. **G**) Main effects of both time [F (2,19) = 7.609, *p* = 0.0037], and sex [F (1,19) = 6.909, *p* = 0.0165] were found in the CeA. Post hocs on the main effect of time showed Fos activity during LA was significantly higher as compared to the EA (*p* = 0.0041) and Rev (*p* = 0.0199) timepoints. **H**) Post hocs on the main effect of time showed the LA timepoint was significantly higher as compared to the EA (*p* = 0.0041) and Rev (*p* = 0.0199) timepoints. The 2-way ANOVA of changes in Fos+ within the BLA only revealed a main effect of time [F (2,18) = 13.65, *p* = 0.0002], and post hocs on the main effect shows differences in EA (*p* = 0.0044) and LA from Rev (*p* = 0.0002). **I**) The LHb, however, had both a significant main effect of time [F (2,23) = 3.776, *p* = 0.0382], and time x sex interaction [F (2,23) = 7.585, *p* = 0.003]. The post hocs revealed that EA and LA males (*p* = 0.0417), as well as LA and Rev males *p* = 0.0004), were significantly different from one another, but no females timepoints differed. **J)** A heatmap representation of the mean Fos+ cell activity of each brain region at each timepoint for both males and females. **K)** A schematic showing all brain regions examined for *C-fos* activity during the targeted helping task.

*Significant change from HCC, p<0.05
#Significant difference from EA, p<0.05
&Significant difference from LA, p<0.05

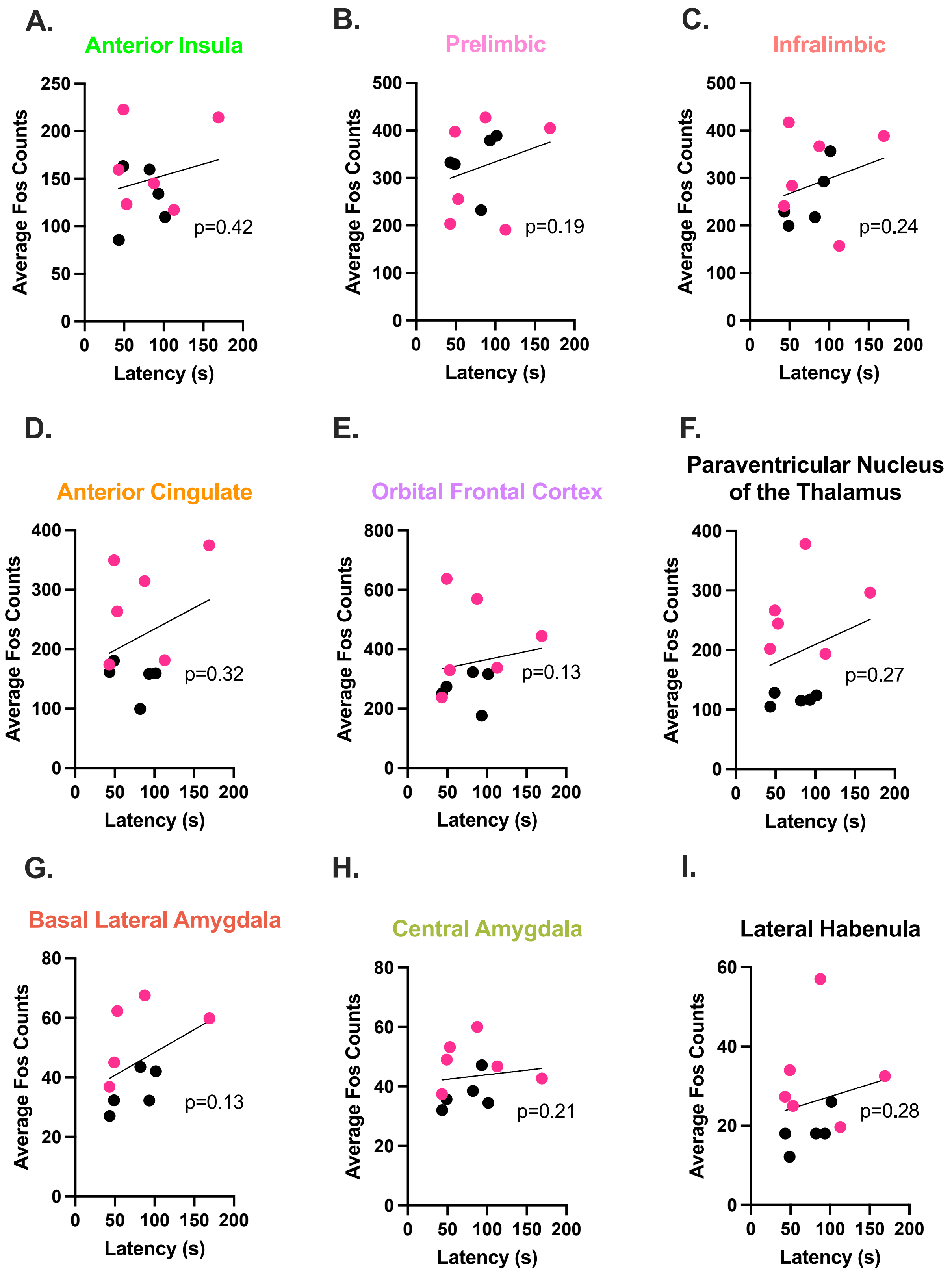


**Supplemental Figure 4. No Significant Correlation Found between Release Latency and Fos Activity during Reversal** Correlational analyses were performed between reversal latency and the corresponding mean Fos+ cell counts for male (data points in black) and female (red) Observers in the following regions of interest: anterior insula (**A**), prelimbic (**B**), infralimbic (**C**), anterior cingulate (**D**), and orbitofrontal (**E**) cortices, paraventricular nucleus of the thalamus (PVT) (**F**), central amygdala (**G**), basolateral amygdala (**H**), and lateral habenula (**I**). Unlike the acquisition timepoints (**Figures 7 & 8**), no significant correlations were seen during reversal.
